# DCMax - A novel Delayed C_max_ Drug Delivery System for Improved Chronotherapy

**DOI:** 10.64898/2026.09.24.753692

**Authors:** Kushagr Punyani, Julie Bondgaard Loehde, Matthias Manne Knopp, Soren Gregersen, Sahil Gupta, Daniel Bar-Shalom

**Affiliations:** Prolevi Bio AB, Lund, Sweden; Steno Diabetes Center Aarhus, Aarhus University Hospital, Denmark; Department of Biomedicine, Health, Aarhus University, Denmark; Bioneer A/S, Department of Pharmacy, DK-2100 Copenhagen, Denmark; Department of Clinical Medicine, Health, Aarhus University, Denmark; Department of Pharmacy, DK-2100 Copenhagen, Denmark

**Keywords:** Controlled release, Chronotherapy, Erosion-based drug delivery, Triiodothyronine, Caffeine, Hypothyroidism

## Abstract

This study evaluated the Delayed C^max^ Drug Delivery System (DCMax), an erosion-based controlled-release formulation designed to delay peak plasma concentration (Cmax) and sustain drug release. The tablet matrix contained hydroxypropyl methylcellulose, hydroxypropyl methylcellulose acetate succinate, and polyethylene oxide. *In-vitr*o dissolution and erosion studies confirmed the delayed release profile. *In-vivo* performance was assessed in six healthy volunteers by measuring salivary caffeine concentrations after administration of immediate-release caffeine tablets and DCMax tablets. DCMax induced a delayed C^max^ of 5 hours compared with 1 hour for the immediate-release formulation, while peak concentrations were lower compared to immediate release. These findings demonstrate that erosion-based drug delivery can achieve delayed release, with potential application in chronotherapy for endocrine disorders.

**Highlights:**

- Erosion-based controlled release tablet may enable chronotherapy dosing
- Chronotherapeutically-relevant release rate achieved for molecules including thyroid hormone T3 and caffeine in-vitro
- Drug release rate largely independent of molecule type when the drug load is ≤20%
- Pharmacokinetic modelling predicts API release profile matching circadian rhythm
- Clinical study shows delayed Cmax and reduced peak during release

## Introduction

The circadian rhythm regulates a wide range of physiological processes, and its disruption can significantly influence the development and progression of various diseases. The circadian clock plays a critical role in several endocrinological processes, such as hepatic glucose production, glucose utilization in skeletal muscles and TSH stimulation for T3 and T4 release [1]. Optimizing the treatment of certain diseases may be possible by aligning therapies with daily circadian biology [1]. Specifically, chronotherapy aims to synchronize treatment with the circadian rhythm [2]. This approach can be implemented through various strategies, such as improving the sleep-wake cycle or timing medication dosing, that can potentially decrease side effects and enhance treatment efficacy [2,3].

Hypothyroidism is a prevalent endocrinological disorder, with overt hypothyroidism affecting up to 5.3% of the population in Europe [4]. However, latest data in 16 major markets in 2024 [5], indicated the combined prevalence to be as high as 7.3%, with substantial variation between countries (range 0.61-12.8%). Its prevalence varies between countries partly due to factors such as iodine deficiency. Women are more frequently affected by hypothyroidism compared to men [4]. The current standard treatment involves oral administration of thyroxine (T4), which is subsequently converted into the active hormone triiodothyronine (T3) [6]. However, a subset of patients continues to experience persistent symptoms despite putatively adequate T4 therapy. An alternative approach involves direct treatment with active T3, but its short half-life necessitates multiple daily doses, thereby complicating its use [6,7]. In addition, increased serum T3 concentrations are associated with adverse effects such as tachycardia [6]. While evidence supporting combined treatment with T4 and T3 over T4 alone remains limited, the American Thyroid Association and European Thyroid Association recently issued a consensus statement addressing this combination treatment and need for controlled release T3 formulations [7].

Pharmaceutical formulations designed for controlled release of active ingredients may improve treatment outcomes, particularly for conditions like hypothyroidism, where replicating the natural nocturnal peak of thyroid hormone levels could be beneficial [8].

The term “Immediate Release” (IR) is used in pharmacopoeias, such as the United States Pharmacopeia (USP) and the European Pharmacopoeia (Ph. Eur.), to describe oral dosage forms that, in dissolution testing, release at least 85% of the active pharmaceutical ingredient (API) content within 30 min. In practical terms, no deliberate attempt is made to influence the rate of release of the drug from the dosage form.

Several terms are used to describe drug delivery systems, particularly oral systems, in which the release of the drug from the dosage unit does not follow an immediate-release profile. These include “Controlled Release,” “Modified Release,” “Depot Systems,” “Delayed Release,” “Extended Release,” and “Long Acting.” There is often confusion and overlap in the use of these terms. Here, “Controlled Release” (CR) is used as a general term, and “Delayed C^max^” (DCMax) refers to the specific approach presented in this article.

Controlled-release technology encompasses a variety of mechanisms, such as erosion and diffusion, to alter or extend the release profile of a drug over time. Compared with IR formulations, extended-release formulations often require less frequent dosing and maintain steadier drug concentrations in the bloodstream [9]. The formulation investigated in this study relies on a surface-erosion-based release mechanism, in which the active substance is gradually released as the tablet matrix erodes, and is therefore referred to as the Delayed C^max^ Drug Delivery System (DCMax).

The DCMax tablet matrix comprises HydroxyPropyl MethylCellulose (HPMC) and/or HydroxyPropyl MethylCellulose Acetate Succinate (HPMC-AS) and PolyEthylene Oxide (PEO). PEO is synthesized by polymerization of ethylene oxide. HPMC is a cellulose derivative in which some free hydroxyl groups in cellulose are replaced with hydroxypropyl and methyl groups [10]. In HPMC-AS, additional hydroxyl groups are substituted with acetate and succinate groups.

These materials are typically used as coatings to delay drug release into the digestive tract by reducing water permeability. In the DCMax tablet investigated in this study, the active ingredient is predominantly released through surface erosion. However, *in-vivo* performance is expected to depend on the physiological functions of the gastrointestinal tract, which are challenging to replicate accurately *in-vitro* or in animal models [11].

This study presents *in-vitro* and clinical data suggesting potential effectiveness of the controlled-release DCMax tablet. *In-vitro* data were obtained from dissolution and erosion studies. Clinical data were obtained using a non-pharmaceutical, commercially available, food-grade test product, containing caffeine. Salivary caffeine concentrations were compared following administration of an IR tablet and the DCMax tablet.

## Materials and Methods

### Materials

For the *in-vitro* dissolution studies, the DCMax tablet formulation contained T3 as the Active Pharmaceutical Ingredient (API), while melatonin, ibuprofen, caffeine, or carvedilol were used as the model proxy molecules. T3 was sourced from Peptido GmbH (Ph. Eur. R1-CEP 2006-249 CAS: 55-06-1,), Melatonin (Sigma Aldrich, CAS # 73-31-4), ibuprofen (Ph. Eur., Fagron A/S,19A28-B10-368735), caffeine anhydrous (BP, Ph. Eur., pharmaceutical grade, PanReac AppliChem), or carvedilol (JP, Ph. Eur., Tokyo Chemical Industry Co., Ltd. respectively. Cytomel containing 20 μg T3 tablets (Orifarm Generics AB, ATC code H03AA02) were used as immediate-release comparators .

For the clinical study, the controlled-release DCMax tablet contained 100 mg caffeine anhydrous (BP, Ph. Eur., pharmaceutical grade, PanReac AppliChem). The matrix components included 360 mg polyethylene oxide (PEO), specifically SENTRY™ POLYOX™ WSR N80-LEO NF Grade, supplied by DuPont (CAS 25322-68-3), and 40 mg HydroxyPropyl MethylCellulose Acetate Succinate (HPMC-AS) under the trade name AQOAT®, sourced from Harke Pharma (Shin-Etsu No. 71045127-03-01, CAS 71138-97-1, Batch 0123294). The tablet coating consisted of EUDRAGIT®, a methacrylic acid-methyl methacrylate copolymer (1:1) conforming to Ph. Eur. and NF standards (CAS 25086-15-1).

The controlled-release tablets (DCMax-Caffeine) were manufactured by Bioneer A/S (Denmark) using direct compression under non-GMP but clean conditions on behalf of Prolevi Bio AB while Cofi-Tabs containing 100 mg caffeine (Vitabalans OY) were bought off counter.

A minimum of 1 mL saliva was collected for human studies using plain plastic tubes with a minimum volume capacity of 1.5 mL, and handling procedures followed the study protocol.

### Dissolution Studies and Analysis

Dissolution studies were conducted using a USP Apparatus 2 (paddle method) with 250 mL vessels (Erweka DT 70, Heusenstamm, Germany). DCMax tablets and immediate-release (IR) tablets containing the active pharmaceutical ingredients (APIs) were tested in triplicate for each formulation (n = 3-6). The dissolution medium consisted of 0.1 M phosphate buffer at pH 6.8, with a total volume of 250mL prepared and preheated to 37°C.

Dissolution of the PEO tablets (n = 3) were studied in USP2 vessels with 125 mL 0.01 M HCl (pH 2.0) at **150 rpm** for 0.5 hours simulating the humans stomach followed by a medium change to 250 mL 0.1 M phosphate buffer pH 6.8 at **50 rpm** for an additional 23.5 hours simulating the human intestine. Samples were taken at 15, 30, 60, 90, 120, 150, 180, 210, 240, 300, 360 and 1440 min and analyzed for drug content using HPLC-UV. After sampling, 0.5 mL fresh medium was added to maintain a constant volume.

Collected samples were transferred directly into High-Performance Liquid Chromatography (HPLC) vials for analysis. Quantification of dissolved caffeine or ibuprofen was performed using HPLC with ultraviolet detection (HPLC-UV). An Ultimate 3000 HPLC system (Dionex, Sunnyvale, USA) equipped with a reverse-phase Kinetex 100A XB-C18 column (4.6 × 100 mm, 5 µm; Phenomenex, Værløse, Denmark) was used. The mobile phase consisted of 0.1% acetic acid (solvent A) and 0.1% acetic acid in acetonitrile (solvent B), with the following gradient elution program: 50% B at 0 min, 80% B at 2 min, 100% B at 3 min, 55% B at 7 min, and 50% B at 10 min. The flow rate was 1.0 mL/min, and detection was carried out at 230 nm. Standard curves for APIs, ranging from 1 to 250 μg/mL, were prepared in triplicate to ensure accuracy and reproducibility.

The mechanism of release of API from the DCMax tablet was investigated via surface dissolution imaging of DCMax tablet (150 mg containing 2 mg ibuprofen) using an SDi2 flow-through dissolution system (Pion Inc., UK). Briefly the tablet was held in position in the flow-cell with metal wires and flushed with a release media (phosphate buffer, pH 6.5 containing 12 g 2mm glass beads, sourced from Pion Inc., at 6.16 mL/min flowrate). The dissolved drug was visualized at 255/280/300/320 nm, while the dissolution of the physical material and hydration were assessed at 520 nm, at an image capture frequency of 50 mHz.

### Clinical Study and Sample Collection

This exploratory, non-randomized, non-blinded crossover study was conducted at Steno Diabetes Center Aarhus, in compliance with Danish regulations and ethical guidelines (approved by the Ethics Committee in Central Denmark Region (1-10-72-161-23)). Samples were analysed by Bioneer A/S. Six healthy adults participated, each serving as their own control by receiving both a controlled-release DCMax tablet and an IR caffeine tablet (Cofi-Tabs) on separate occasions, with a washout period of 1-2 weeks. Both tablets contained 100 mg caffeine.

Participants aged 18-75 years with a body mass index (BMI) of 18.5-35 kg/m² were screened using predefined criteria. Exclusion criteria included pregnancy, cardiovascular or metabolic disorders, caffeine sensitivity, and substance use. Informed consent was obtained following full disclosure of study procedures.

Dietary restrictions included abstaining from caffeine-containing foods and beverages for 24 hours and fasting for 12 hours before administration. Water intake was restricted for two hours before dosing, except for 240 mL carbonated water taken with the tablet. No water was consumed within 10 min of saliva sampling. A caffeine-free meal was provided six hours post-administration, with no further caffeine intake until the final sample collection at 24 hours.

Saliva samples were collected at −60 and 0 min before dosing, and at 0.5, 1, 2, 3, 4, 6-, 7-, 8-, and 24-hours post-dosing.

### Clinical Sample Analysis

Caffeine concentrations in saliva were measured using HPLC-UV as described above. Samples were stored at 4°C and shipped immediately to Bioneer A/S for analysis. All samples were processed and analysed within five to seven days of collection and destroyed after completion of analysis in compliance with ethical standards.

### Contributions

*In-vitro* erosion and dissolution studies were performed and analysed by Bioneer A/S and sponsored by Prolevi Bio AB. For the clinical study, Prolevi Bio AB (the sponsor) designed and prepared the study protocol. The clinical study was conducted at Steno Diabetes Centre Aarhus. Saliva sample analysis was performed by Bioneer A/S and Prolevi Bio AB. Statistical analysis of clinical study data was performed by Bioneer A/S. The manuscript was drafted collaboratively by authors from Prolevi Bio AB, Bioneer A/S and Steno Diabetes Center Aarhus.

## Results G Discussions

### Optimized DCMax tablets demonstrate erosion based uniform API release

To determine the rate of release of the active substance, DCMax tablets (200 mg with 50 μg API, T3) or IR tablets (20 mg with 20 μg API, T3) were dissolved in a USP Apparatus 2 (paddle method) with 2 L of 0.1 M phosphate buffer at pH 6.8 and 37°C, modelling the gastro-duodenal environment. Diluted samples were taken every 30-60 min for up to 5 hours and at 24 hours (data not shown) and analysed using UV-HPLC for quantification of the API or model proxy molecule. As shown in Figure 1a, the IR tablets exhibited an immediate-release profile with approximately 100% of the molecule released within the first hour. The DCMax tablets showed a gradual release of the API or model proxy molecule reaching over 80 % over the first five hours, approaching 100% over 6 hours.

**Figure 1.**
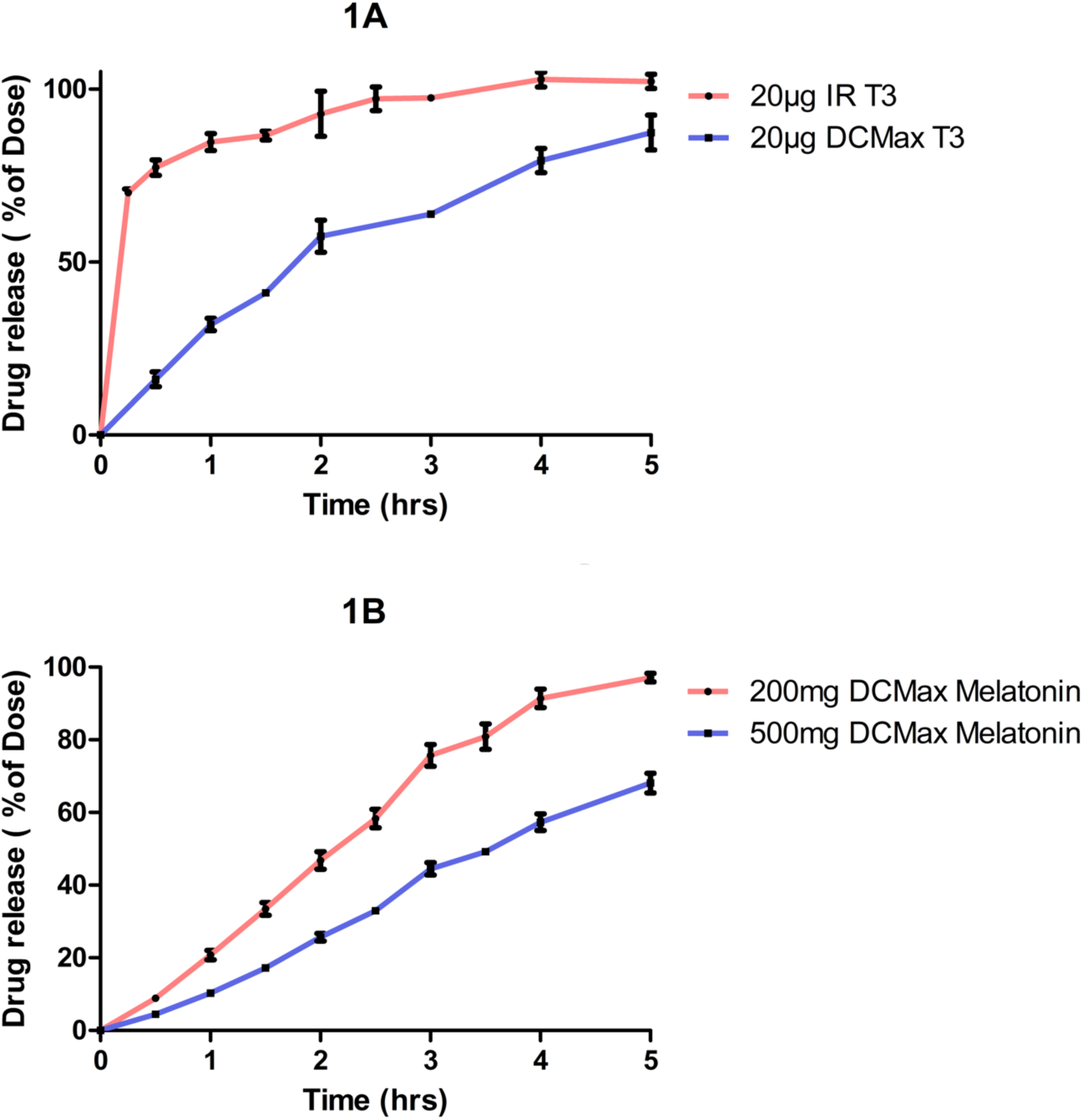
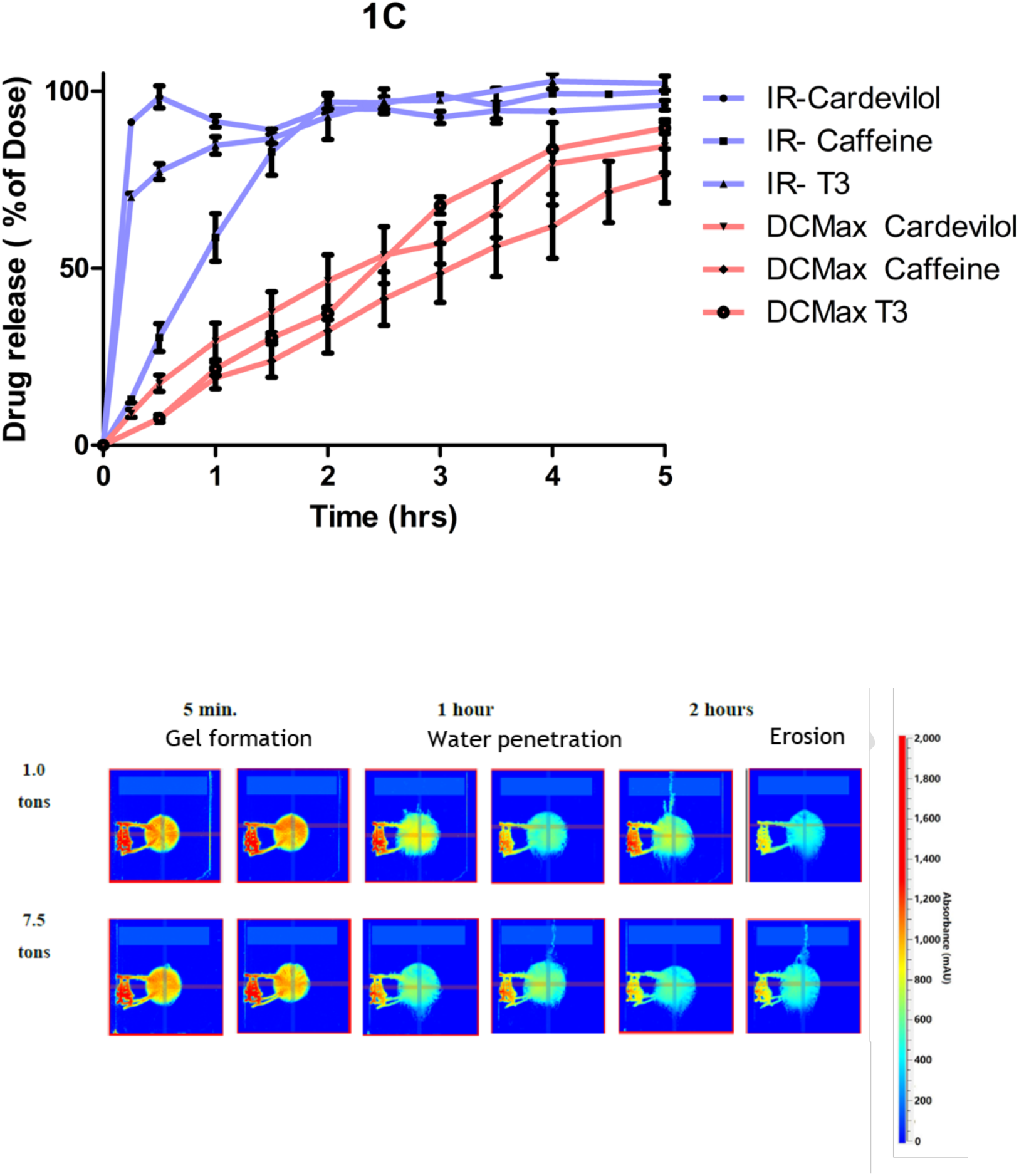
Dissolution profiles, size effects, and imaging of erosion-based release from DCMax tablets. Dissolution profile of IR tablets (blue) and DCMax tablets (red) containing 50 μg T3, 100mg Caffeine, 5µg Melatonin performed in a USP Apparatus 2 (paddle method). (a) IR tablets (20µg T3) released ∼100% of API within 1 h, while DCMax tablets released gradually, approaching 100% over 5 *h*. (b) Effect of DCMax tablet mass (200 mg vs. 500 mg) on the release of 5 mg melatonin as a model proxy molecule with same concentration. Both sizes produced similar release profiles, although the larger tablet (500mg) exhibited a slower release rate. (c) Dissolution of IR tablets (blue) vs DCMax tablets (red) containing T3, caffeine and carvedilol. All DCMax tablets showed similar release profiles *independent* of the API of choice. (d) Surface dissolution imaging of two DCMax tablets (150 mg), compressed by 1 one or 7,5 tons, each containing 2 mg ibuprofen, used to visualize the release mechanism. The gel layer is shown in blue and the intact tablet core in red.

To determine the impact of size or geometry on drug release from the DCMax tablet, dissolution studies were performed with tablets weighing 500 mg and 200 mg (Fig. 1b) using 5 mg melatonin as the model proxy molecule. Melatonin was selected due to its similarity to T3 as a hormone, its relevance for chronotherapeutic release, and its low weight-to-volume ratio compared with tablet geometry. This also enabled assessment of whether different APIs would show similar release rates. The profiles were similar for both the sizes; however, the larger tablet demonstrated a slower release.

To investigate the effect of API on the release profiles from DCMax tablets, dissolution studies were performed on DCMax and IR tablets containing T3, melatonin and carvedilol. As shown in Fig. 1c, all DCMax tablets showed a release profile, independent of the API. To further investigate the release mechanism (diffusion or erosion), UV imaging was conducted with ibuprofen as a model proxy molecule. Upon hydration, encompassing gel formation and water penetration phases, marked by a reduction in surface channels or cracks, the dosage unit gradually eroded, forming only a thin release layer on the surface. This process indicated that the active molecule was released primarily through erosion (Fig. 1d). The tons relate to the pressure applied when compressing the tablets. Beyond one ton there is no change in release because the penetration of water into the matrix reached a plateau (or, conversely, the “tightness” did)

### Predicted Pharmacokinetic Profiles

To predict the pharmacokinetic profile of DCMax tablets, the release profiles of melatonin tablets weighing 200 mg and 500 mg, each containing 5 mg melatonin, were convoluted following the approach described elsewhere [12]. As shown in Figure 2a, the release profiles and areas under the curve for both tablet sizes were similar. However, the larger tablet showed a reduced C^max^ and a longer dissolution peak.

**Figure 2.**
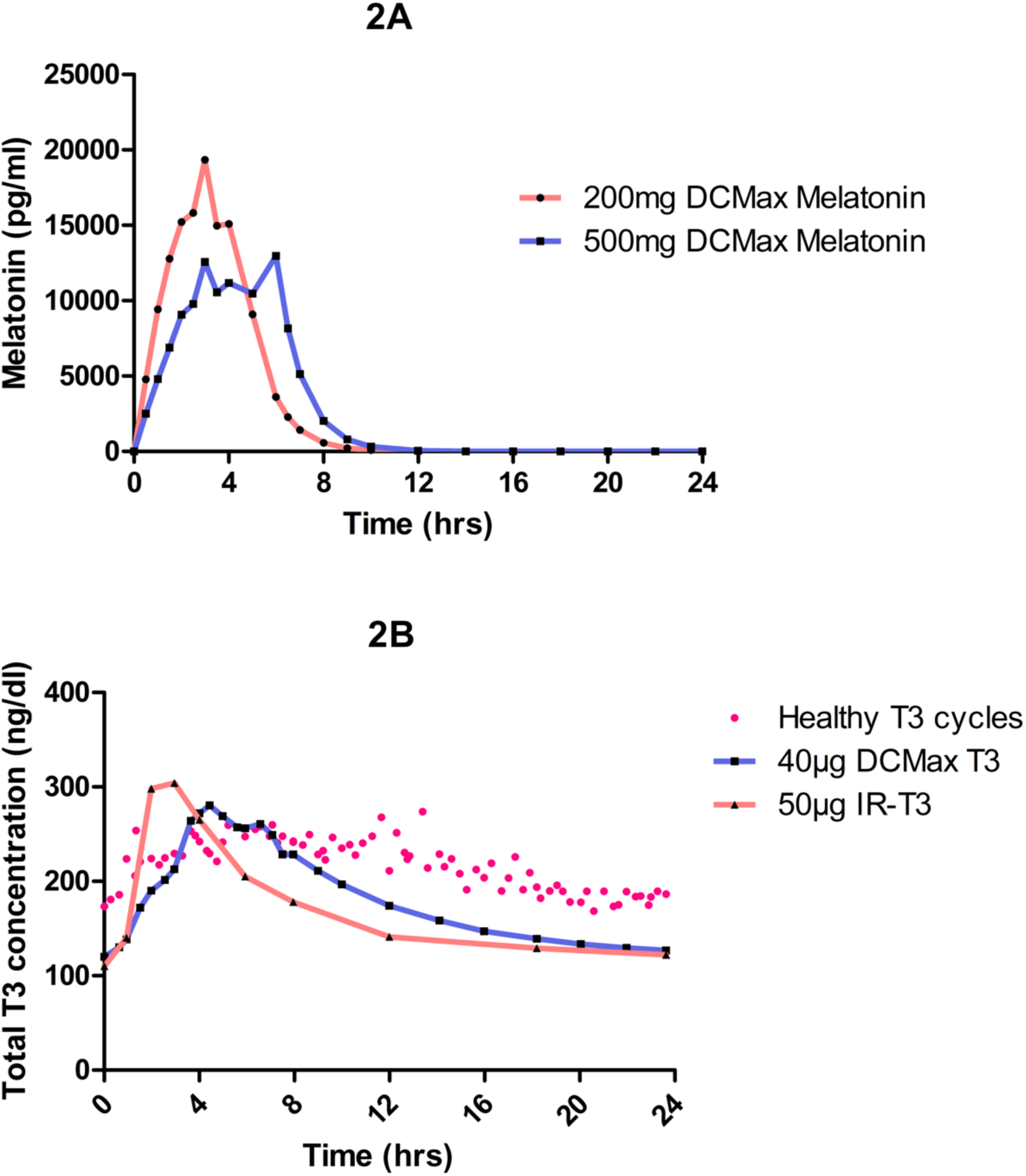
Predicted plasma concentration-time profiles for DCMax formulations and immediate-release comparators. (a) Predicted plasma concentrations of DCMax melatonin tablets weighing 200 mg (red) or 500 mg (blue), each containing 5 mg melatonin. Both tablet sizes produced similar exposure (area under the curve), although the 500 mg tablet showed a lower C^max^ and a slower release. (b) Comparison of predicted plasma concentrations of DCMax tablets containing 40 μg T3 (IR-T3, blue) and commercial IR Cytomel tablets containing 50 μg T3, red. An endogenous T3 plasma concentration profile over 24 h following the circadian rhythm is shown in red dots (adapted from [13]). DCMax produced a slower release with a 4-5h delay in T^max^ and maintained T3 concentrations within the therapeutic window for a longer duration than Cytomel.

The same model was applied to evaluate DCMax and Cytomel, a commercially available IR formulation of T3. As shown in Figure 2b, DCMax tablet containing 40 μg T3 demonstrated a slower release profile, with 3-4 hours delay in peak concentration, compared with IR formulations containing 50 μg T3, which returned to baseline approximately 12 hours after administration. The predicted T3 profile for DCMax remained within the therapeutic window and closely matched physiological T3 plasma concentrations that follow the circadian rhythm [13]. A 50 μg T3 DCMax tablet (previously published data [10] showed similar results to 40 μg T3 DCMax tablet (rate of drug release, Suppl Fig.1); however, the 40 μg dose was selected based on the postulated improved efficacy, as its C^max^ remained within the therapeutic window.

## Clinical Performance

### Participant Demographics and Compliance

Six healthy individuals (five females, one male) participated in the study comparing the clinical release profiles of IR and DCMax tablets, each containing 100 mg caffeine. Baseline characteristics were mean age 48.5 ± 12.8 years, height 174.4 ± 6.8 cm, weight 72.5 ± 11.3 kg, and body mass index (BMI) 23.8 ± 2.9 kg/m². All participants completed the study without dropouts. One participant experienced headache, assigned to abstaining from caffeine prior to the study day and was treated with paracetamol.

### Caffeine Concentration in Saliva

Caffeine concentrations in saliva were measured for up to 24 h after administration (Fig. 3). IR caffeine tablets produced a steep increase in caffeine concentration shortly after dosing. In contrast, DCMax produced a more gradual increase, significantly delaying the mean time to maximum concentration (T^max^) from 1.83 ± 0.17 h (IR) to 6.17 ± 0.54 h (DCMax) (P = 0.004). Consistent with the expected reduction in peak concentrations, DCMax showed a significantly lower C^max^ than IR tablets (1.43 ± 0.09 μg/mL vs 2.11 ± 0.06 μg/mL, P = 0.003). DCMax also appeared to produce a lower area under the curve (AUC, µg mL^-1^ hr) for caffeine (17.73 ± 1.84 vs 21.69 ± 1.78 for IR)(P = 0.051).

**Figure 3.**
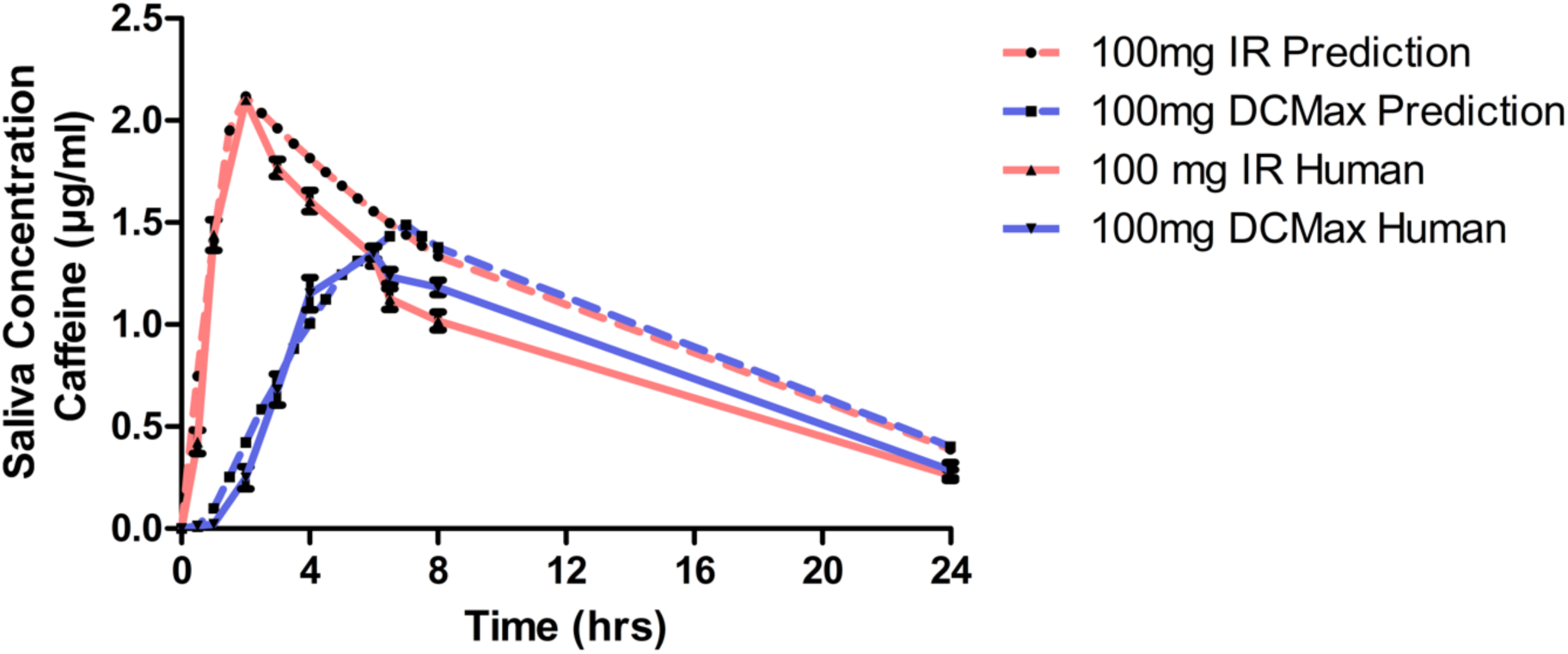
Caffeine concentration-time profiles in saliva following administration of IR and DCMax tablets. Mean (±SD?) caffeine concentrations in saliva over 24 h for study participants (n=C) after administration of a 100 mg IR caffeine tablet (red) or a 100 mg controlled-release DCMax tablet (blue). No saliva samples were collected between 8 h and 24 h post-dose.

## Discussion

The simplicity of DCMax tablets lies in their composition, which is readily scalable using standard manufacturing techniques such as dry granulation, roller compaction, and hot melt extrusion [10,14]. The formulation uses two or three excipients and achieves high content uniformity (data not shown). We aimed to test of efficacy of convoluted data i.e. PK prediction from in vitro dissolution assays to in human tested data. Furthermore, we wanted to develop a controlled-release tablets with delayed drug release that mimic with natural hormone rhythms, i.e., chronotherapy. The hypothesis was that better mimicking of natural hormone levels, or timing drug dosing to coincide with physiological peaks and troughs, would translate into improved therapeutic efficacy.

The data from the in vitro studies in this article demonstrate that, irrespective of the active molecule used (ibuprofen, melatonin, T3, caffeine) in the DCMax tablet, the rate of release is constant when the drug load is ≤20%. At this dosing, release is mediated primarily through erosion, ensuring homogeneous delivery. To predict human pharmacokinetic profiles from i*n-vitro* dissolution data, we employed a published convolution method [12]. To validate these predictions, published human Cytomel data were compared with model predictions showing a 1:1 match between observed and predicted PK profiles (Suppl. Fig 2).

Since the primary objective was to assess the technological and commercial viability of T3, caffeine was selected as a proxy compound. Caffeine, a nutraceutical with well-documented human safety and toxicity data, provided a suitable proof-of-concept candidate and its dissolution profile in the DCMax tablet was comparable to T3 at equivalent tablet geometry and drug loading. No active ingredient in the DCMax tablet is chemically modified, and all are stable ingredients with well documented human toxicity.

The clinical study showed a delayed Tmax and decreased Cmax with the DCmax tablet compared to IR tablet, however, there also appeared to be a trend to decreased AUC with the DCmax tablet. As there were no samples obtained between 8 h to 24 h, the caffeine release within this time period is unknown and warrants further investigation.

Animal models, while valuable for studying human pathologies, have limitations in replicating human peripheral or local thyroid hormone responses. Rodents display multiphasic sleep during the light phase, whereas humans have a consolidated nocturnal sleep period [15–17]. As circadian rhythm is directly linked to the sleep-wake cycle [13], rodent models are not ideal for studying endocrine hormone regulation in the context of circadian rhythm. This could lead to misleading results in therapeutic screening. Moreover, maintaining the geometry of erosion-based controlled-release tablets in small animal models is challenging.

The results in this study offer proof of release patterns with the DCMax technology and support the accuracy of convolution models in predicting human responses from *in-vitro* data.

## Conclusion

This study evaluated the effectiveness of a controlled-release formulation, here called DCMax. The tablets exhibit a “hill shaped”, delayed Cmax rate of API release, achieving complete release within 6 h as shown by *In-vitro* dissolution testing, convolution-based PK predictions*. In-vivo* caffeine clinical trial data showed a lower maximum concentration (Cmax) and a longer time to maximum concentration (Tmax) with the DCmax tablets.

## Disclosures

Daniel and Sahil have been employed at and hold financial securities in Prolevi Bio AB. Kushagr is a board member and consultant at, and holds financial securities in Prolevi Bio AB. Matthias, Julie and Sören have been contracted by Prolevi Bio AB, as described in the article.

## Author Contributions

The authors are listed alphabetically under CRediT contributions. Conceptualization: Daniel, Kushagr, Sahil. Data curation: Julie, Matthias, Sahil. Formal analysis: Daniel, Matthias, Sahil, Sören.

Funding acquisition

Kushagr, Sahil. Writing - original draft: Julie (clinical study), Kushagr, Sahil. Writing - review and editing: All authors.

**Supplementary Fig.1:**
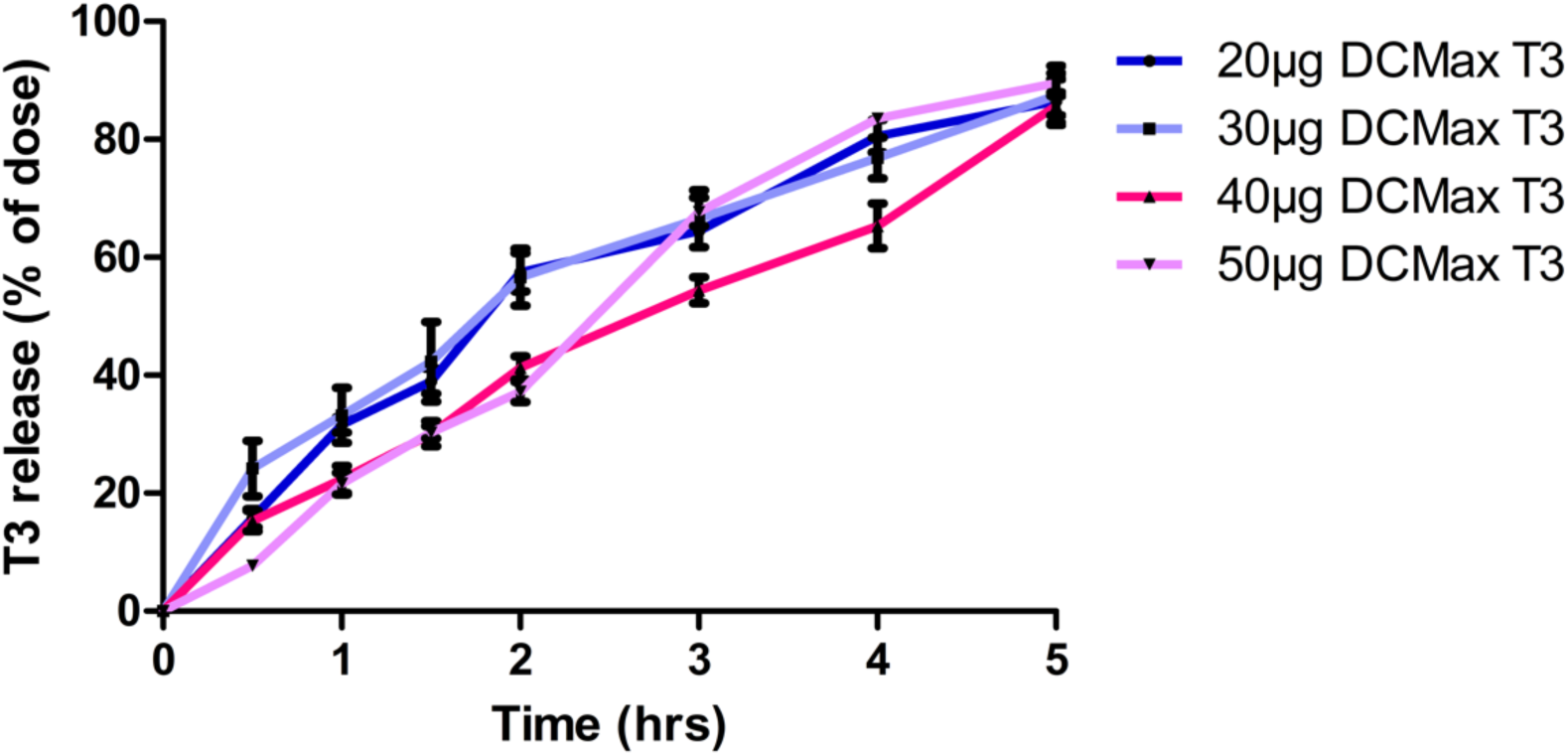
Rate of release from dissolution profiles of 20µg, 30 µg, 40 µg, 50µg T3 DCMax tablets manufactured with dry granulation analyzed by ELISA demonstrate similar rate of release irrespective of API dose.

**Supplementary Fig. 2:**
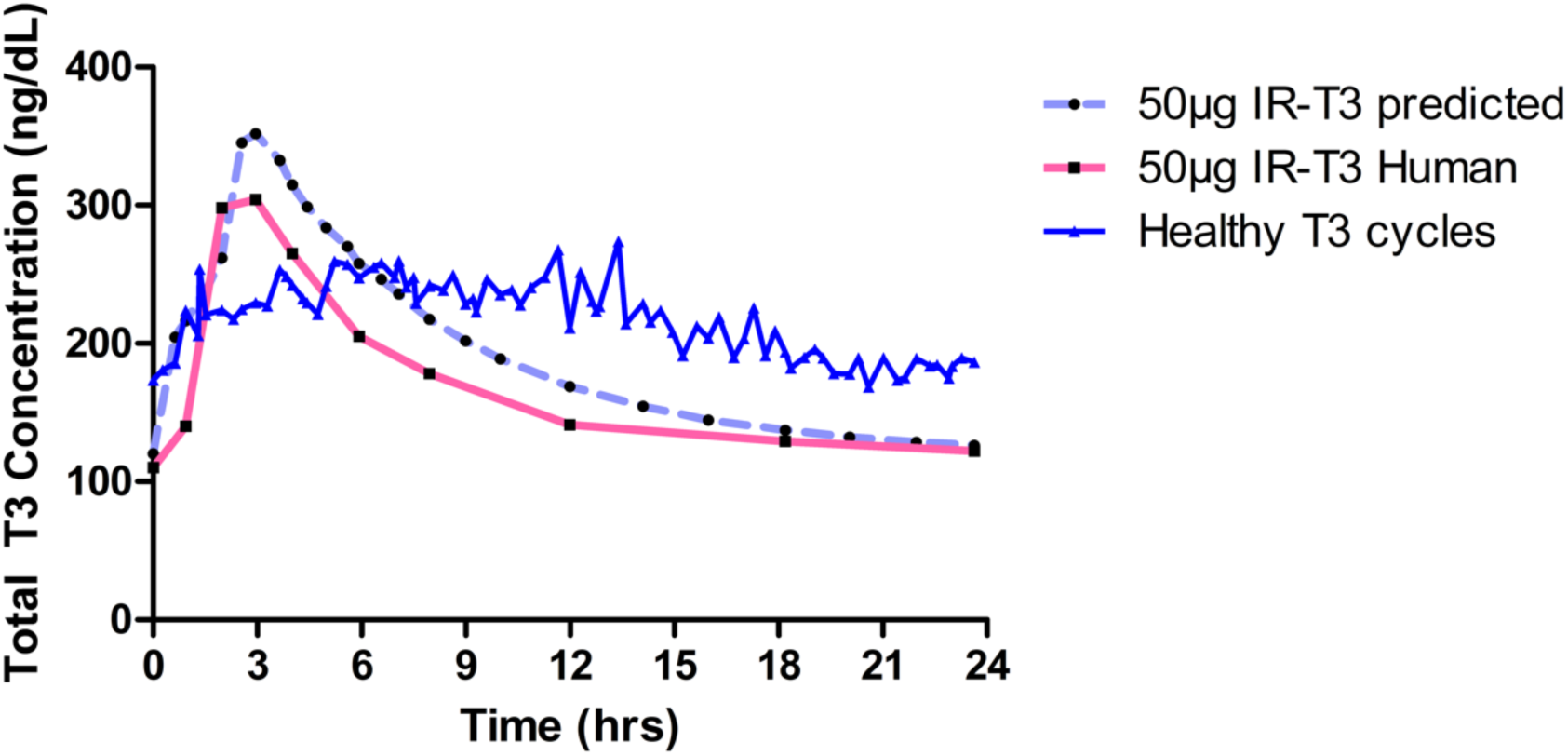
Pharmacokinetic profiles from in human data of 50 µg of IR-T3 was compared to 50µg DCMax convoluted data generated from in vitro dissolution assays (Prolevi’s data, Synthonics PZL-T3, ITL Pharma BCT-303).

## Notes

### Competing Interest Statement

DBS and Sahil Gupta have been employed at and hold financial securities in Prolevi Bio AB. KP is a board member and consultant at, and holds financial securities in Prolevi Bio AB. MMK, JBL and
Soren Gregersen have been contracted by Prolevi Bio AB, as described in the article.

### Funding

- VINNOVA: 2021-03873

