## Supplementary figures and images for "DCMax - A novel Delayed C_max_ Drug Delivery System for Improved Chronotherapy"

### Supplementary Figure 1

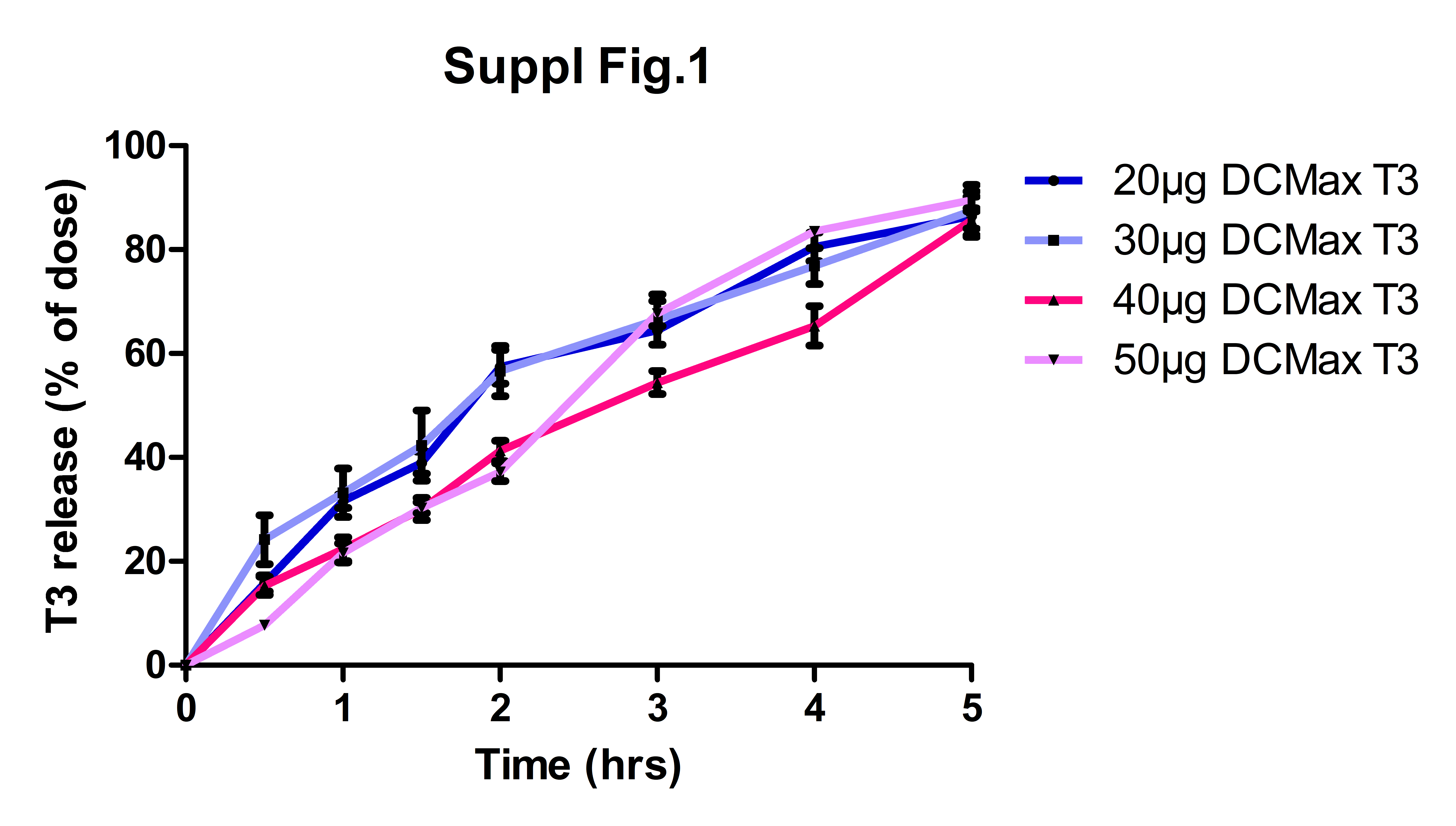

### Supplementary Figure 2

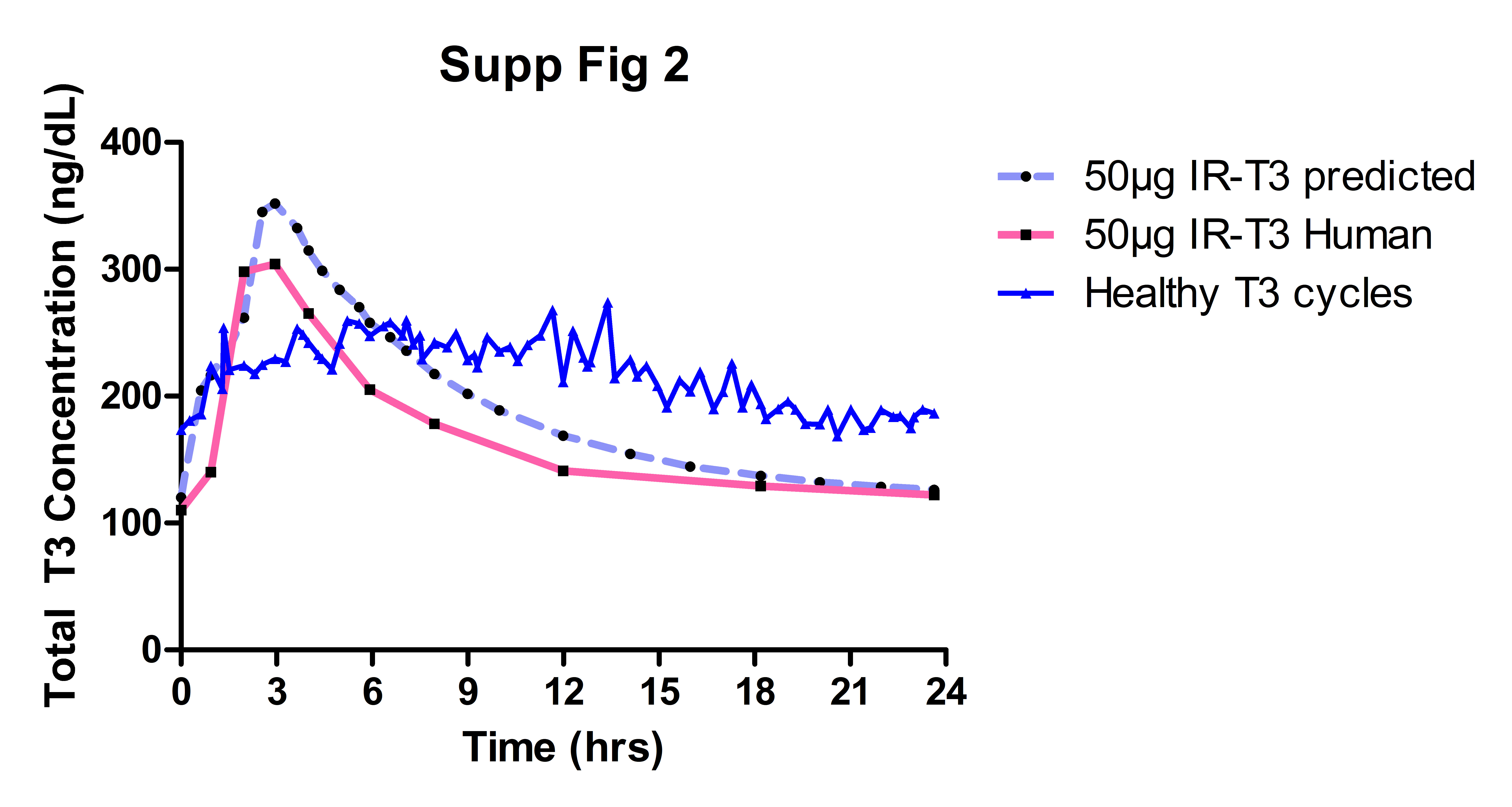
